# MINTsC learns multi-way chromatin interactions from single cell high throughput chromatin conformation data

**DOI:** 10.1101/2024.04.08.588641

**Authors:** Kwangmoon Park, Tianchuan Gao, Jingwen Yan, Sündüz Keleş

## Abstract

A number of foundational analysis methods have emerged for single cell high throughput chromatin conformation (scHi-C) datasets capturing 3D organizations of genomes with pairwise measurements at the single cell or nuclei resolution; however, these datasets are currently under-utilized. The canonical analyses of scHi-C data encompass, beyond standard cell type identification, inference of chromosomal structures and pairwise interactions. However, multi-way chromatin interactions among genomic elements are entirely overlooked. We introduce **MINTsC** to learn **M**ulti-way **INT**eractions from **s**ingle cell Hi-**C**. MINTsC builds on a dirichlet-multinomial spline model and yields multi-way interaction scores by aggregating pairwise interactions across cells of a context and summarizing them using order statistics of pairwise test statistics. MINTsC yields well-calibrated p-values for controlling the false discovery rate. Evaluation of MINTsC with scHi-C datasets from cell lines and complex tissues using multiple external genomic and epigenomic datasets support multi-way interactions inferred by MINTsC. Application of MINTsC to scHi-C data from human prefrontal cortex revealed multi-way chromatin interactions with biological implications of gene regulation by multiple enhancers. Most notably, MINTsC-inferred multi-way interactions demonstrate its potential for probing molecular QTL and association studies for epistatic SNP effects by significantly reducing the multiple-testing burden. MINTsC is publicly available at https://github.com/keleslab/mintsc.

## 1 Introduction

Advances in high-throughput single-nucleus mapping of 3D genome organization via pairwise contacts between proximal loci now enable genome-wide characterization of loops, TADs, A/B compartments, and subcompartments at single-nucleus resolution. Yet these datasets have not been systematically leveraged to resolve multi-way chromatin interactions (simultaneous contacts among multiple loci within the same nucleus). Identifying such interactions is crucial for assessing the epistatic potential of risk variants from genome-wide association studies (GWAS) and molecular quantitative trait loci (QTL) studies. Disease-linked genes are often regulated by multiple distal enhancers spread over large genomic distances, e.g., nested enhancer cooperativity of the oncogene MYC was dissected by multiplexed CRISPRi screening ^1^. Genome-wide association studies have shown that more than 90% of disease-associated variants are noncoding variants^2, 3^, and can be dispersed over large distances. Furthermore, while single enhancer variants may exert modest effects individually, combinations can interact to substantially influence traits and disease.

A suite of experimental innovations now measure multi-way chromatin contacts: GAM ^4^, ChIA-drop ^5^, SPRITE ^6, 7^, Tri-C, multi-contact 4C ^8^, COLA ^9^, and Pore-C ^10^. These platforms established feasibility mainly in cell lines and mESCs, but Hi-C and scHi-C remain the most widely generated and used for complex tissues. Notably, the NIH BRAIN Initiative has produced extensive high-quality scHi-C from human and mouse brains. Yet analyses of scHi-C have largely focused on pairwise contacts, often adapting bulk Hi-C methods (with snapHi-C ^11, 12^ as a notable exception). This underutilizes scHi-C for elucidating how multiple enhancers cooperate over long genomic distances to regulate target genes and confer disease risk. Long-read sequencing-based scNanoHi-C ^13^ demonstrated that multi-way contacts can be captured in single cells, but cellular heterogeneity and experimental noise make them inconsistently observable across cells. scMicro-C ^14^ likewise revealed dynamic variability even in relatively homogeneous cell lines. These observations underscore the need for a principled statistical framework that models the underlying multi-way chromatin interaction structure and reconstructs it from noisy, heterogeneous single-cell data.

In this study, we introduce MINTsC, the first method to learn “**M**ulti-way Chromatin **INT**eractions from scHi-**C**”. We cast multi-way chromatin interaction discovery as clique detection in a multilayer network: each layer is a cell, nodes are genomic loci, and edges denote locus-locus contacts (**Fig**. 1a). MINTsC models pairwise contacts with an empirical Bayes spline null that accounts for the well-known genomic distance bias of scHi-C assays, and then derives a clique-level test statistic with an analytic null distribution. Multiple testing is controlled via the Benjamini-Hochberg (BH)^15^ procedure to call multi-way chromatin interactions at a target false discovery rate (FDR). Because the clique statistic aggregates pairwise signals, MINTsC also applies targeted pre- and post-processing filters to minimize spurious multi-way chromatin interaction calls.

**Figure 1.**
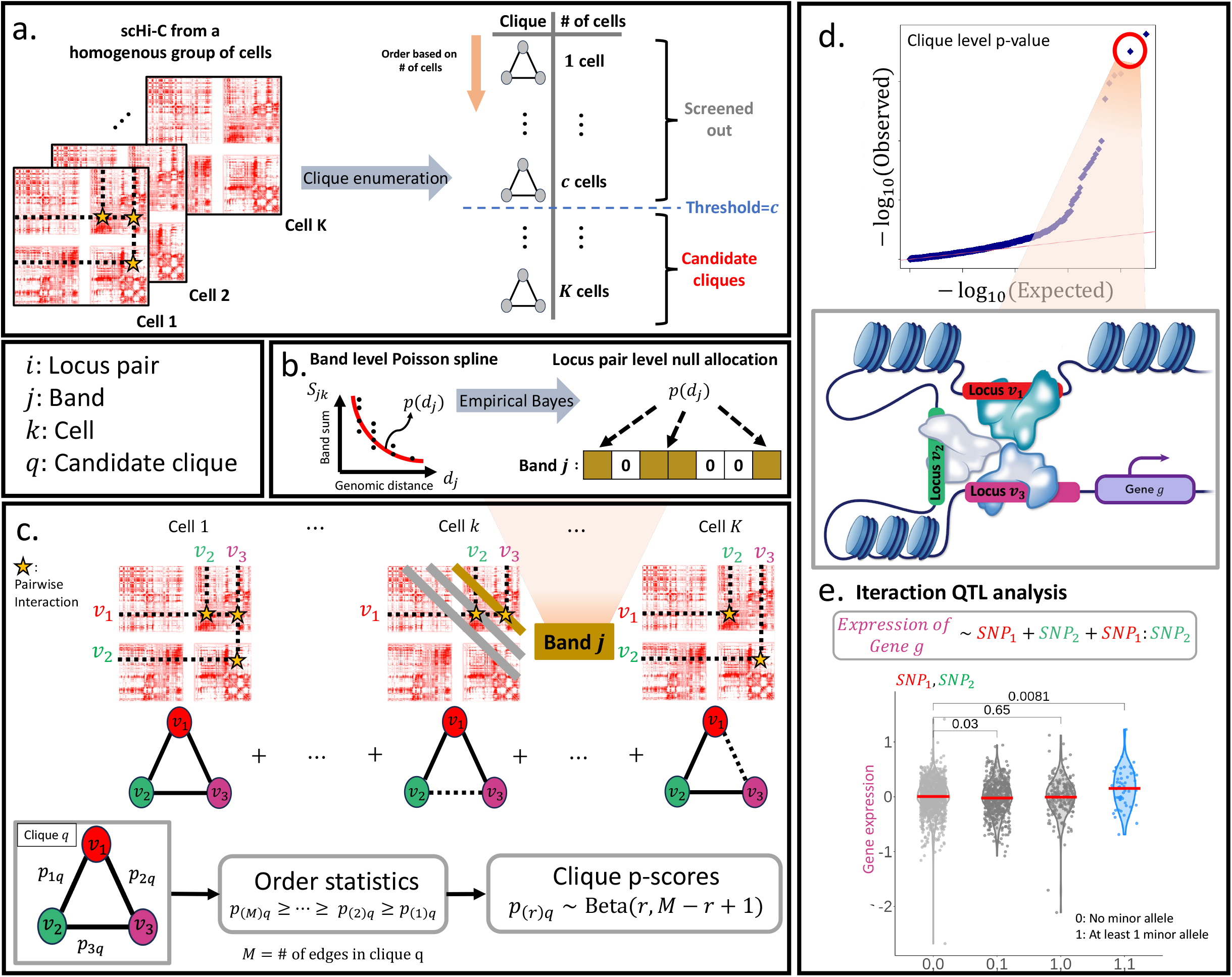
A schematic overview of MINTsC for learning multi-way interactions from scHi-C data. **a**. MINTsC first employs a clique search algorithm on the scHi-C contact matrices of a homogeneous group of cells, e.g., a cell type, and enumerate how many times each clique appears across cells. Then, a cell number threshold is applied to the set of cliques searched to define *candidate cliques*. **b**. MINTsC fits a locus-pair level multinomial model to scHi-C contact counts by a multinomial aggregation and poisson approximation strategy at the band level. Contact counts *y*_*ijk*_ of locus pairs, *i* ∈ [*N*_*j*_], across bands, *j* ∈ [*J*], of a cell *k* ∈ [*K*] follow a multinomial distribution. Band-level aggregated counts for each band *j* within cell *k*, 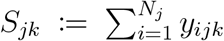, follows a poisson distribution with a rate parameter that depends on the band-level count probability which is modeled as natural cubic spline function *p*_*j*_ = *p*(*d*_*j*_) of genomic distance *d*_*j*_ between the locus pairs in the band. Estimated band level probabilities, *p*_*j*_, are distributed across the locus-pairs within a band with under a null Dirichlet prior to the multinomial counts (**Methods**). **c**. MINTsC multi-way interaction scores (i.e., clique-level test statistics) are based on the order statistics of pairwise interaction test statistics, e.g., p-values. Square matrices denote scHi-C contact matrices of *K* cells for a fixed chromosome. The starred entries depict the pairwise contacts among three genomic loci (i.e., nodes) *v*_1_, *v*_2_, *v*_3_, displayed as an undirected graph in the bottom. The gray or gold bars along the off diagonals denote bands. MINTsC aggregates the pairwise contact counts within a given candidate clique across the cells of the cell type, and generates clique-level statistic, which are clique p-score *p*_(*r*)*q*_ (**Eqn**. 7 in **Methods**) and clique z-score 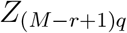 (**Eqn**. 8 in **Methods**). **d**. Q-Q plot of the − log_10_(clique p-values), with *x*- and *y*-axes denoting − log_10_ transformed uniform random variables and the observed clique p-values (**Eqn**. 9 in **Methods**), respectively. **e**. Illustration of a multi-way interaction among genomic loci harboring two enhancers and a gene promoter. MINTsC multi-way interactions can be probed for epistatic SNP effects in molecular QTL studies or GWAS.

Evaluations of MINTsC with external multi-way chromatin interaction data, imaging-based 3D genome data, and data from methylation assays support the multi-way interactions identified by MINTsC. Applied to scMicro-C data, MINTsC calls remain consistent with inferred 3D genomic distances despite the dynamic nature of multi-way chromatin contacts representing the contributions of individual enhancers to the regulation of genes. Compared with a baseline derived from state-of-the-art scHi-C loop calling, MINTsC identifies more multi-way chromatin interactions with markedly better FDR control in human prefrontal cortex and synthetic data. Strikingly, interrogation of MINTsC-inferred multi-way interactions from human prefrontal cortex in an eQTL study of human cortex reveals epistatic SNP effects for genes linked to amyloid-*β* pathology and synaptic plasticity.

## 2 Results

### 2.1 Overview of MINTsC

Bulk Hi-C’s aggregate nature cannot resolve multi-way chromatin interactions. In contrast, scHi-C enables a graph formulation in which multi-way contacts correspond to cliques in a multilayer network: each layer is a cell, nodes are genomic loci (bins; typically 10 kb-1 Mb depending on coverage), and edges encode locus-locus contacts ^16^ (**Fig**. 1a). Despite ligation limits and sparsity, scHi-C yields abundant cliques (**Fig**. S1), confirming its potential to infer lower-order multi-way chromatin interactions (empirically order 3-6), even if very high-order events remain challenging.

We developed MINTsC to learn multi-way chromatin interactions from scHi-C by testing cliques for significance. First, MINTsC fits an empirical-Bayes spline null model for pairwise contacts that accounts for genomic-distance bias, yielding bias-adjusted pairwise test statistics (**Fig**. 1b; Methods). Next, within a homogeneous context (e.g., a cell type or treatment), MINTsC aggregates pairwise evidence across cells to form a clique-level statistic (“clique p-score”) using order statistics of the pairwise p-values, and derives an analytic null distribution for this statistic (**Fig**. 1c; Methods). Under the homogeneity assumption (Supplementary Note), the clique score reflects the strength of the corresponding clique in the population-level contact matrix. Because the statistic is built from distance-adjusted pairwise tests, it is stable to the distribution of intra-clique genomic distances (**Fig**. S2). This yields well-calibrated *clique p-values* based on parametric null distribution of the clique p-scores for FDR control (**Fig**. 1d, **Method**) and facilitates downstream analysis, e.g., interaction eQTLs, by drastically reducing the multiple testing burden. For example, the expression variability of a gene with promoter within locus *v*_3_ can be studied for contributions of the interactions between SNPs (e.g., SNP_1_ in *v*_1_ and SNP_2_ in *v*_2_) that reside in loci *v*_1_ and *v*_2_ rather than exhaustively over all combinations of gene’s *cis* SNPs (**Fig**. 1e).

Because multi-way calls aggregate pairwise contacts, MINTsC includes pre- and post-filtering to guard against spurious multi-way chromatin interactions. A pre-filter defines candidate cliques as those fully observed in at least a minimum number of cells, protecting against spurious cliques formed by merging pairwise interactions from different groups of cells (**Fig**. 1a; Methods). An optional post-filter applies a statistical co-occurrence test on clique edges to further eliminate artifacts (Supplementary Note).

### 2.2 Validation of MINTsC multi-way chromatin interactions for same-cell colocalization

Since MINTsC relies on aggregation of pairwise contacts, we first evaluated fidelity of MINTsC-identified multi-way interactions at the single-cell resolution. Specifically, by leveraging orthogonal 3D spatial distance measurements and targeted simulations, we assessed whether inferred multi-way interactions reflect same-cell co-localization. In all of the applications, we grouped the candidate cliques evaluated by MINTsC into two based on MINTsC clique p-values resulting from the parametric Beta distribution of the clique p-scores (**Methods**). Clique p-values were then subjected to FDR thresholding^15^ and labeled as *multiway chromatin interactions*.

#### (i) 3D spatial distance–based validation with train-test split on GM12878 scMicro-C

We applied MINTsC to scMicro-C contact matrices^14^ of 10kb resolution from GM12878 cells, fitting on one group of cells (48 training cells) and evaluating on the remaining cells (48 test cells). Pre-filtering step of MINTsC generated 43,469 three-way candidate cliques and MINTsC applied to training set identified 904 multi-way chromatin interactions (at FDR of 0.1). Using the 3D genome coordinates constructed for the test scMicro-C cells with the Dip-C algorithm^17^, we computed, for each candidate clique in every test cell, the maximum pairwise 3D distance (Euclidean) among the clique loci. Across the individual test cells, cliques identified as multi-way chromatin interaction by MINTsC showed significantly smaller maximum pairwise 3D distances than other cliques, supporting same-cell colocalization of the interacting loci (Fig. 2a). Notably, for nearly all cells, within clique maximum pairwise 3D distances of MINTsC-multiway chromatin interactions were below 3.5 particle radii (∼240 nm),which is the distance cutoff used in scMicro-C to define pairwise chromatin interactions. Next, to obtain an estimate of empirical FDR, we applied MINTsC to the whole set of cells and quantified the proportion of cells each clique had a maximum pairwise 3D distance below 3.5 particle radii. This revealed that MINTsC-multi-way chromatin interactions co-localized in a significantly larger proportion of cells compared to cliques that were not significant by MINTsC (FDR of 0.1, **Fig**. 2b). We show that the association based on scMicro-C still hold after matching the cliques with similar total pairwise interaction strength, further demonstrating that MINTsC scores are indeed signatures of multi-way chromatin interactions (**Fig**. S3a,b). When cliques with maximum pairwise 3D distance less than 3.5 particle radii in at least 10% of the cells were considered as true multi-way interactions, MINTsC yielded an empirical FDR of 8% (FDR of 10% when the requirement on the proportion of cells was raised to 50%).

**Figure 2.**
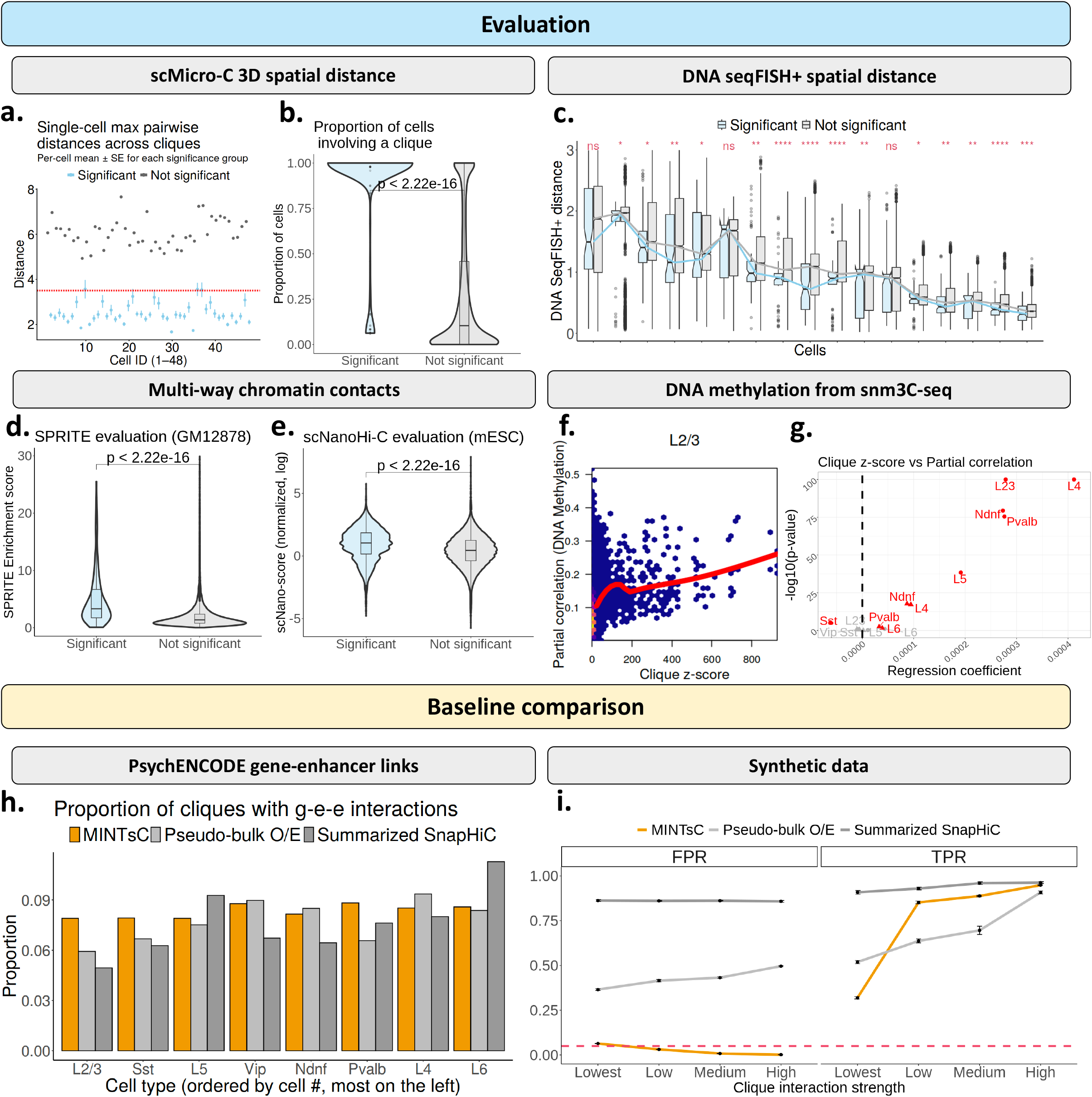
MINTsC identifies multi-way interactions supported by a multitude of external data and with biological implications in gene regulation. **a**. Per-cell mean and two-step standard deviation of the maximum pairwise 3D spatial distance within each clique, stratified by significance group. The red dashed line marks the 3.5 particle radii cutoff suggested by Wu *et al*.^14^ **b**. Proportion of cells where a clique’s maximum 3D distance is below 3.5 particle radii (∼240 nm) ^14^, comparing significant and non-significant MINTsC cliques in GM12878 scMicro-C contact matrices^14^. **c**. Comparison of within-clique DNA seqFISH+ distances between significant and insignificant MINTsC candidate cliques for the top 20% of cells with the highest number of candidate cliques (as indicated by the red vertical line in **Fig**. S4a. The x-axis indexes individual cells. Blue and gray lines denote the trend of DNA seqFISH+ distances across cells for significant and insignificant MINTsC cliques, respectively. **d**. Comparison of SPRITE enrichment scores ^6^ between the significant MINTsC cliques (FDR*<*0.01) and the others for GM12878 ^43^. **e**. Similar comparison for MINTsC mESC multi-way interactions (FDR*<*0.05) using normalized scNanoHi-C ^13^ concatemer counts. **e**. Comparison of clique z-scores (*x*-axis) with average absolute partial correlations of locus pairs within each clique based on single cell DNA methylation (*y*-axis) for L2/3 cells (**Fig**. S11 for other cell types). **f**. − log_10_(p-values) (*y*-axis) of the estimated regression coefficients (*x*-axis) from regressing the MINTsC clique z-scores on the partial correlation summary of DNA methylation. The cases with BH FDR*<*0.05 are labelled as significant. **h**. Proportion of gene harboring cliques with the full *g* − *e*_1_ − *e*_2_ based on PsychENCODE gene-enhancer interactions. Cell types are ordered based on the number of cells from left to right. **i**. Comparison of MINTsC with baseline approaches using synthetic data (**Methods** for details). FPR and TPR for each method are quantified under different signal scenarios for multi-way interactions (*x*-axis) (Red line : *y* = 0.05).

#### (ii) Cross-technology validation in mouse cortex and hippocampus

We next utilized MINTsC to infer multi-way chromatin interactions from mouse cortex and hippocampus scHi-C (Dip-C) data ^18^. The contact maps from Dip-C were binned at 25Kb resolution to match the resolution of the publicly available mouse brain DNA seqFISH+ data ^19^ for evaluation. The DNA seqFISH+ dataset harbored spatial coordinates of targeted 1.5Mb genomic region of each chromosome in individual cells and enabled calculation of spatial distances between loci pairs.

We considered the set of candidate cliques with loci in the profiled genomic regions of the DNA SeqFISH+ data and asked whether, within individual cells, MINTsC-identified multi-way chromatin interactions had smaller maximum pairwise 3D genome distance compared to rest of the cliques. In the top 20% (17/81) of cells with the most profiled cliques (cells with ≥ 6, 000 candidate cliques), MINTsC-identified multi-way interactions had significantly smaller within-clique distances than the remainder (Fig. 2c, p-value *<* 0.01). With a relaxed threshold (cells with ≥ 2, 000 candidate cliques), 71% of the 62 cells showed the same pattern (p-value *<* 0.001). Furthermore, pooling data across all cells further supported this observation: MINTsC-identified interactions in the scHi-C (Dip-C) data corresponded closely to spatial proximity patterns in the DNA seqFISH+ dataset (**Fig**. S4b). We show that the association based on DNA seq FISH+ still hold after matching the cliques with similar total pairwise interaction strength (**Fig**. S3c,d). Compared to the within-assay validation in (i) with scMicro-C, the comparatively reduced separation observed here likely reflects additional cross-modality and between-experiment variability inherent to the DNA seqFISH+ comparison, rather than a limitation of either platform. Lastly, to evaluate the false discovery rates of the method applied to Dip-C data, we leveraged a proximity matrix construction method ^19^ for converting DNA seqFISH+ imaging data to pairwise interaction data. Specifically, for each individual cell in the DNA seqFISH+ dataset, all the candidate cliques with maximum within clique DNA-seqFISH+ distance less than 150 nm ^19^ were labelled as true multi-way interactions. Using this gold standard, MINTsC false positive rate did not exceed 3% at the individual cell level.

#### (iii) Simulation-based evaluation of MINTsC for same-cell co-localization

Although MINTsC infers multi-way chromatin contacts by aggregating pairwise interactions, held-out and per-cell 3D distance evaluations support the co-localization of the identified interactions within cells and their robustness to cellular heterogeneity. We next evaluated this observation with a simulation experiment. We simulated scHi-C contact counts under a multinomial model parameterized from the Lee *et al*. dataset (chromosome 19; 5,582 10 kb bins; L2/3 neurons) (see **Methods** for details on data generation). We spiked in two types of of three-way cliques into the scHi-C contact matrices: (i) triads whose loci co-occurred within individual cells (true cliques) and (ii) triads formed by two pairs co-occurring in one cell group and the remaining pair in another (spurious cliques) (**Fig**. S5a; Methods). Applying MINTsC at an FDR of 0.05 while varying pairwise interaction strengths, the empirical FDR remained well controlled at 5% across settings (**Fig**. S5b). This experiment, specifically designed to lead to spurious aggregation of pairwise contacts as multi-way interactions, supports that the MINTsC framework is robust against aggregation of pairwise interactions originating from different groups of cells.

### 2.3 SPRITE and scNanoHi-C validation of MINTsC multi-way chromatin interactions with both low and high resolution scHi-C data from human cell lines and mouse embryonic stem cells

After evaluating MINTsC for same cell co-localization, we turned our attention to validation of applications to low and high resolution scHi-C data. We first evaluated MINTsC’s ability to infer multi-way chromatin interactions supported by multiple external datasets using GM12878 ^20^ and mESCs (Serum/Lif) ^21^ at resolutions of 500Kb and 1Mb, respectively (**Tab**. S1, **Tab**. S2 for details). As in the Section 2.2 scMicro-C and Dip-C applications, we grouped the cliques from scHi-C into two, based on MINTsC clique p-values. Cliques with adjusted^15^ clique p-values *<* 0.01 were labelled as *multi-way interactions*. For evaluation, we utilized the enrichment scores of the SPRITE multi-way interactions introduced in Quinodoz *et al*. ^6^. These scores aim to denoise raw SPRITE counts by comparing observed counts of a multi-way interaction against counts of randomly constructed multi-way interactions of the same size while matching the genomic distances within the random multi-way interactions and the observed one. Higher enrichment scores are interpreted as strong evidence for multi-way interactions^6^. For both the GM12878 and mESC datasets, for which SPRITE data was available, cliques labelled as multi-way interactions by MINTsC exhibited significantly larger enrichment scores compared to cliques that were not significant (**Fig**. 2d,S6a; Wilcoxon rank-sum test p-values *<* 0.01). Further investigation with clique size stratification revealed that significant cliques from MINTsC exhibit a larger enrichment score mode than that of the insignificant cliques for both GM12878 and mESC (**Fig**. S6b,c). In addition to the alignment of MINTsC with SPRITE, MINTsC inferred multi-way interactions in mESCs are also similarly supported by the GAM triplet score^4^-based evaluation (**Fig**. S7). Furthermore, comparisons of MINTsC *clique z-scores* (**Methods**) against clique-level summarizations of pairwise SPRITE and external bulk Hi-C data revealed strong positive associations (**Fig**. S8, Supplementary Note).

Next, we turned our attention to higher resolution (10Kb) mESC (2i/Lif) scHi-C data and leveraged the recent long-read based single cell high throughput chromatin conformation (scNanoHi-C) data of mESC (2i/Lif)^13^ for evaluating MINTsC multi-way chromatin interactions. scNanoHi-C assay generated long sequencing reads that contained multiple DNA monomers, referred to as “concatemers”. Consequently, concatemers that include DNA monomers originating from three or more different genomic locations signify interactions that are multi-way. We investigated the genomic distance and depth normalized concatemer counts of multi-way interactions (**Fig**. 2e) and observed that multi-way interactions inferred from MINTsC have significantly higher scNanoHi-C counts across clique sizes compared to cliques that are not deemed as multi-way interactions (one-sided Wilcoxon rank-sum test p-value *<* 2 × 10^−16^). We also show that these associations based on SPRITE and scNanoHi-C still hold after matching cliques with similar total pairwise interaction strength (**Fig**. S9). Further stratification of the cliques based on their within clique maximum genomic distance revealed that cliques with maximum within clique genomic distance less than 200Kb are well supported by the scNanoHi-C data (**Fig**. S10), suggesting an empirical genomic distance constraint for post-processing the significant cliques from MINTsC.

Collectively, these results suggest that MINTsC can identify multi-way interactions in both high and low resolution scHi-C data from cell lines. In what follows, we showcase the applicability of MINTsC with more recent human and mouse brain tissue scHi-C datasets.

### 2.4 Evaluation of MINTsC with high-resolution scHi-C data from complex human and mouse brain tissues

We applied MINTsC to infer multi-way interactions in human prefrontal cortex neuronal cell types (excitatory: L2/3, L4, L5, L6 and inhibitory: Nndf, Pvalb, Sst, Vip) using scHi-C data from sn-m3C-seq assay which simultaneously profiles chromatin conformation capture and methylation at the single cell level ^22^. Data for each cell type was binned at 10Kb resolution to generate contact matrices for each cell. While gold standard multi-way chromatin interactions for this system are not available via other technologies, we designed an approach to assess whether the resulting MINTsC clique z-score, i.e., the *r*th smallest locus pair level z-value within each clique (Methods), associated with multi-way interactions by leveraging the interplay between methylation and chromatin conformation capture at the single cell level. Specifically, we utilized the well-established observation that DNA methylation level correlations (either positive or negative) are indicative of regulatory relationships among the loci ^23–25^. In fact, recent work established that DNA methylation profiles and chromatin accessibility as measured by ATAC-seq at the single nuclei level validated each other at the cell-subtype level ^26^. Hence, genomic loci within a multi-way interaction can be expected to exhibit a dependency structure in their DNA methylation profiles. To assess such dependence, we leveraged partial correlation coefficient and quantified the level of conditional dependence between pairs of genomic loci given the other loci in the multi-way interactions. For example, for a three way interaction, if any of the pairs exhibit low partial correlation in absolute value given the third locus in the candidate interaction, this would indicate that pairwise dependencies within the candidate multi-way interaction are not supported by the co-methylation patterns. With this line of reasoning, we then compared MINTsC clique z-scores with the average DNA methylation-based partial correlations between all the loci of individual cliques, at clique sizes of 3 and 4 and across all the eight cell types (**Fig**. 2f,S11). This analysis revealed that cliques with larger MINTsC clique z-scores tend to have larger multi-way dependency levels across all clique sizes for each cell type. We further assessed the association between the MINTsC clique z-scores and partial-correlation based multi-way dependencies (slopes of the fitted regression lines, **Fig**. 2g). In this quantification, majority of the comparisons yielded a significant and positive association between the MINTsC clique z-scores and partial-correlation based evidence for multi-way interaction.

Having observed that the MINTsC multi-way chromatin interactions exhibit single-cell level co-localization based on DNA seqFISH+ 3D distances in Section 2.2, we performed a comparative analysis between the MINTsC clique *z*-scores and the DNA seqFISH+ distances summarized at the clique level. We focused on the cell types shared by both datasets (Dip-C^18^ and DNA seqFISH+^19^) and compared the clique *z*-scores with the 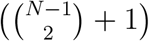 th closest spatial distance between loci pairs within the *N* -way cliques. Specifically, we chose the order number as 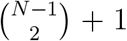 to ensure that all the *N* loci are participating to form a connected graph. This comparison indicated a consistent inverse relationship (high MINTsC z-score with low spatial distance) across the cell types (asctrocyte, microglia, and oligodendrocyte progenitor cells) and clique sizes (3,4) (**Fig**. S12a,S13). A permutation-based evaluation confirmed that loci within MINTsC-inferred multi-way interactions were significantly closer spatially than expected by chance (**Fig**. S12b). Assessment of this inverse association with slopes of the fitted regression lines (**Fig**. S12c) further yielded a significant negative association between the MINTsC clique z-scores and the spatial distances of loci within multi-way interactions for majority of the cell types.

### 2.5 Application of MINTsC to haplotype-resolved Dip-C data

It is worth remarking that the mouse brain scHi-C data from the Dip-C assay^18^ is haplotype-resolved. This allowed us to investigate multi-way interactions in a haplotype-aware manner. Specifically, we deployed MINTsC on single cell contact counts of individual haplotypes. This yielded that MINTsC clique p-values are well calibrated (**Fig**. S14), making it readily applicable to haplotype-resolved scHi-C data. A comparative analysis of the MINTsC results for the two haplotypes revealed that the p-values for most cliques are concordant between the two haplotypes across cell types. Further comparison of the haplotype-aware analysis with the aggregate analysis that sums over contact matrices of the two haplotypes suggested that while aggregate analysis results in some power loss in detecting haplotype-specific multi-way chromatin interactions as expected, it does not lead to inflation of false positives (Supplementary Note, **Tab**. S3).

### 2.6 MINTsC identifies significantly more multi-way interactions with higher reliability compared to baseline approach

To the best of our knowledge, MINTsC is the first approach for inferring multi-way chromatin interactions from scHi-C contact matrices. We compared MINTsC to two baseline approaches that we devised from standard analysis of scHi-C data. Specifically, we considered multiway interactions in the form of cliques from **1)** the significant loop calls of SnapHiC ^11^ (i.e., significant pairwise interactions) and from **2)** strong pairwise interactions with pseudo-bulk O/E (Observed/Expected) ≥ 2 based on cell type specific scHi-C data. We deployed these baseline approaches on the human prefrontal cortex scHi-C data. The first baseline approach did not yield cliques with order larger than 3; hence, we retained the comparisons to cliques of size 3. We thresholded the candidate cliques from MINTsC at FDR 0.05 to obtain significant 3-way interactions.

Next, we employed external PsychENCODE data ^27^ to evaluate the resulting multi-way interactions of MINTsC and the baseline methods. Specifically, we utilized the compiled neuronal cell type gene-enhancer interactions which encompassed one or more interactions per gene. After filtering the cliques of each method so that at least one locus in the clique harbored at least one gene, we quantified how many of the method cliques were of the form *g* − *e*_1_ − *e*_2_ structure (v-structure), where *g* refers to the gene with enhancers *e*_1_ and *e*_2_ in the gene-enhancer link list from the PsychENCODE source. Specifically, for each gene *g* within a locus of a clique *q*, we assigned the clique *q* a score of 1, if the other loci in the clique individually overlapped the PsychENCODE enhancers of the gene *g*. Under this scoring scheme, a clique could have score *s* if it contains *s* number of *g* − *e*_1_ − *e*_2_ v-structures. Next, we quantified proportion of method cliques with scores *s* ≥ 1 (**Fig**. 2h). This calculation was carried out separately for each cell type and yielded that MINTsC outperforms the baseline approaches in terms of capturing the *g* − *e*_1_ − *e*_2_ structures across cell types (**Fig**. S15).

We further conducted baseline comparisons using synthetic scHi-C datasets, mimicking the characteristics of actual scHi-C data (**Fig**. S16a). Specifically, we leveraged the Lee *et al*. ^22^ scHi-C dataset (chromosome 19 with 5,582 10Kb-sized bins, and L2/3 neurons) and multinomial distribution for the scHi-C counts in generating synthetic data for the comparisons. To ensure a more heterogeneous background contact count distribution than accounted for by MINTsC (i.e., not strictly following the MINTsC background), we randomly implanted true (300) and false (2,100) cliques in the estimated background contact probability by varying the signal sizes of the pairwise contacts within true cliques (**Methods** for details on the synthetic data generation). We then evaluated the False/True Positive Rates (FPR/TPR) across the simulation replications. While the TPRs of the considered approaches were comparable, MINTsC exhibited significantly lower FPRs (*<* 0.05) (**Fig**. 2i) across signal sizes of the true cliques, due to its well-calibrated pairwise and multi-way interaction p-values (**Fig**. S16b,c,d and **Fig**. S17). These results underscore MINTsC’s capability to identify multi-way chromatin interactions with more reliable FDR control while maintaining comparable power to natural baseline methods.

### 2.7 Biological implications of MINTsC-inferred multi-way interactions involving genes and multiple enhancers

After establishing that multi-way interactions inferred by MINTsC are supported by external data, we turned our attention to specific multi-way interactions that involve genes and enhancers. For these investigations, we leveraged (i) Activity-By-Contact (ABC) scores ^28^ that infer gene-enhancer interactions based on genomewide profiling of chromatin accessibility and activating histone mark H3K27ac; and (ii) gene-enhancer interactions inferred from multi-modal profiling of single cells for chromatin accessibility and gene expression ^29^.

We considered the mESC^30^ cliques evaluated by MINTsC and leveraged the ABC scores from Narita *et al*. ^31^ to asses whether multi-way interactions with at least one gene are supported by the ABC scores of the involved enhancers. A total of 6,030 cliques harbored genes regulated by multiple enhancers within the same clique, i.e., cliques containing a full *g* −*e*_1_−*e*_2_ −· · ·−*e*_*N*_ structure, where *g* refers to a gene with ABC scores evaluated for enhancers *e*_1_, *e*_2_, ·· ·, *e*_*N*_ . Specifically, we required the cliques to have one locus overlapping with a gene and have the other clique loci each overlap with at least one enhancer for which the gene has a reported ABC score. For each clique satisfying this requirement and each *g* −*e*_1_ −*e*_2_ −· · ·−*e*_*N*_ structure in the clique (6,584 cases in total), we took the average of the ABC scores assigned for all *g* −*e*_*j*_ links, *j* ∈ {1, ···,*N* }, to summarize ABC scores at the clique level. Then, we assessed if the multiway interactions inferred from MINTsC (FDR cutoff of 0.05) are supported by the multi-way gene regulation level, which is represented by the clique level ABC score. This comparison revealed a significant difference in summarized ABC scores of significant and insignificant cliques (**Fig**. S18, one-sided Wilcoxon rank-sum test *p* < 2*e* − 16).

Next, for the Lee *et al*.^22^ human prefrontal cortex scHi-C data analyzed at 10Kb resolution, we conducted a similar analysis for the cliques with a full *g* − *e*_1_ − *e*_2_ − · · · − *e*_*N*_ structure, with the enhancers *e*_*j*_s from the set of prefrontal cortex enhancers identified by an integrative analysis of snATAC-seq and scRNA-seq data (further details in **Fig**. 2 of Li *et al*. ^29^). Specifically, Li *et al*. ^29^ provides a set of *g* − *e*_*j*_ links referred to as positively correlated gene-cCRE connections. We compared the MINTsC − log_10_(p-scores) of the cliques containing a *g* − *e*_1_ − *e*_2_ −· · · − *e*_*N*_ (with positively correlated gene-cCRE connections) against those of randomly selected cliques and observed that cliques supported by the gene-cCRE connections tend to have markedly stronger MINTsC multi-way interaction levels (**Fig**. S19). A consistent result can be observed across cell types based on a more systematic comparison with multiple replicates of the randomly selected cliques (**Fig**. 3a).

**Figure 3.**
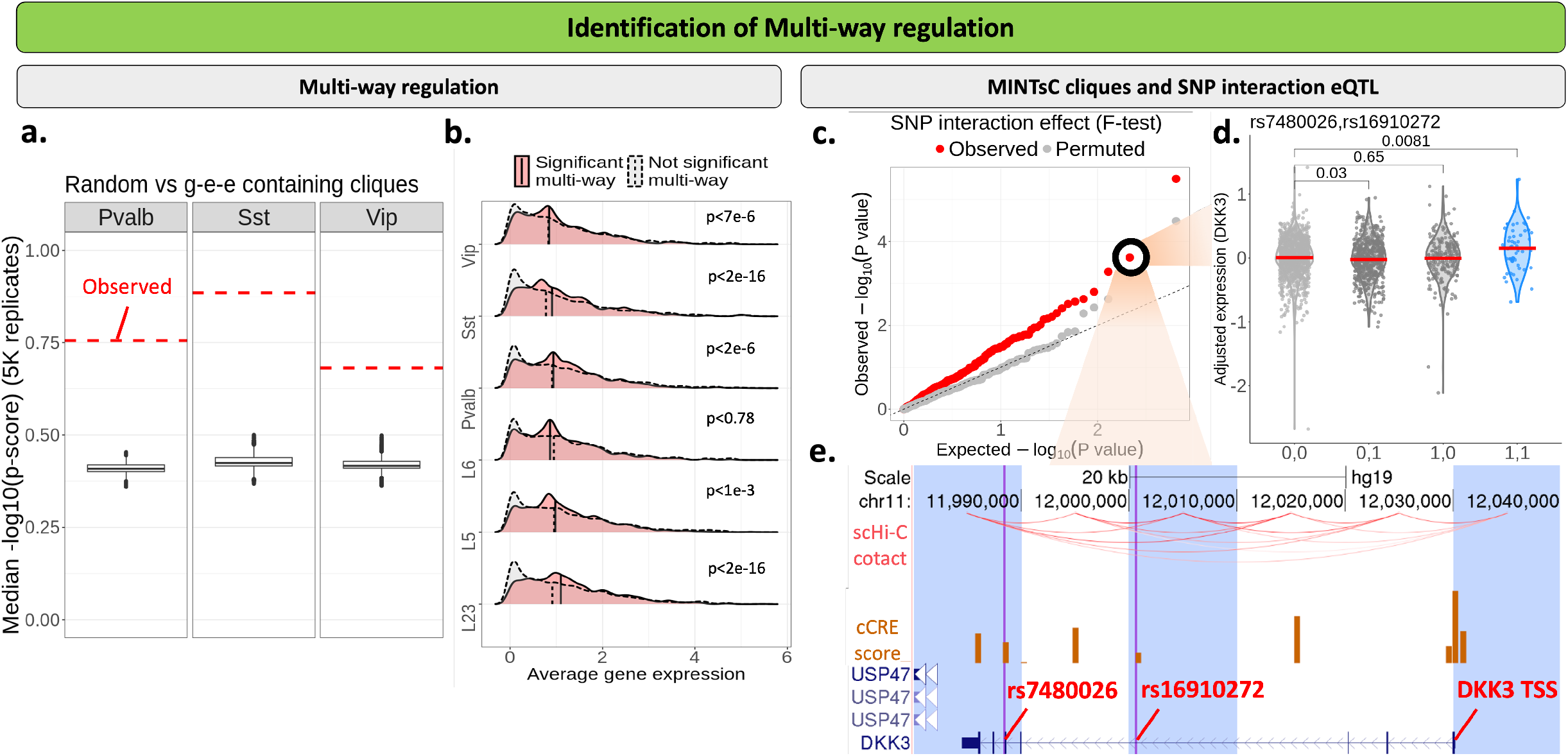
MINTsC identifies multi-way interactions supported by biological implications in gene regulation. **a**. Summarization of the median MINTsC − log_10_(p-scores) of the randomly sampled cliques across 5K replicates. The red line on each panel (cell type) denotes the median − log_10_(p-scores) of the *g* − *e*_1_ − *e*_2_ (from the gene-cCRE list^29^) harboring cliques. **b**. Comparison of average (normalized) gene expression levels among cliques with (FDR*<*0.1) and without significant multi-way interactions across the cell types in human brain prefrontal cortex^22^. p-values above each density plot is based on one-sided Wilcoxon rank-sum test. **c**. QQ-plot displaying F-test − log_10_(p-values) of SNP interaction effects on gene expression from AD ROS/MAP study (red) and that of under random permutation (grey). Each red dot is from a two-way SNP interaction eQTL model, where the gene TSS and SNP locations are identified as a multi-way interaction by MINTsC (Vip cells). **d**. Covariate-adjusted expression levels of DKK3 across samples for combinations of alleles of SNPs rs16910272 and rs7480026. For each SNP, ‘0’ indicates no minor allele, and ‘1’ indicates existence of at least one minor allele. BH adjusted p-values are reported for each comparison. **e**. UCSC Genome Browser display of the DKK3 multi-way interaction. Orange bars depict ATAC-seq signal summarized by cCRE scores^29^ and the red arcs denote the pseudo-bulk scHi-C contacts ^22^ for Vip cells. Purple vertical lines within each blue shaded 10Kb bin denote the SNP locations with significant interaction effect on DKK3 expression.

To probe biological implications of multi-way interactions, we asked whether genes with MINTsC-inferred multi-way interactions exhibited different gene expression patterns than those without multi-way interactions by leveraging the Lee *et al*.^22^ scHi-C data. Utilizing SMART-seq data ^32^ of the same system, we compared expression levels of the genes with and without multi-way interactions for individual cell types. **Fig**. 3b compares the expression distribution of the genes within the two groups and shows that the genes with multi-way interactions on average exhibit significantly higher expression levels throughout most of the cell types (Wilcoxon rank-sum test p-values ≤ 0.003). This agrees well with the findings from the long-read based scHi-C which revealed a similar result for GM12878 cells ^33^.

### 2.8 MINTsC-derived multi-way interactions in human prefrontal cortex enable discovery of potential epistatic SNP effects

To establish the broader biological discovery potential of identifying multi-way interactions, we investigated whether MINTsC could facilitate discovery of epistatic SNP effects in eQTL analysis. Specifically, we focused on pre-frontal cortex multi-way interactions (FDR*<*0.05) involving genes and their enhancers supported by cCRE scores (**Tab**. S4). We leveraged the ROS/MAP study (**Methods**) of Alzheimer’s disease, and assessed whether variation in cerebral cortex expression of genes involved in multi-way interactions inferred by MINTsC were better explained by an eQTL model with SNP-SNP interactions compared to a linear additive model. We specifically investigated the p-values of the ANOVA F-test statistics for the null hypothesis *H*_0_ : *β*_12_ = 0 comparing the full model with a two-way SNP interaction term, i.e., Expression = SNP_1_*β*_1_ + SNP_2_*β*_2_ + (SNP_1_ : SNP_2_)*β*_12_ + error, against the reduced model without the SNP interaction term, i.e., Expression = SNP_1_*β*_1_ + SNP_2_*β*_2_ + error, of each (Gene, SNP_1_, SNP_2_) tuple within a MINTsC multi-way interaction supported by non-zero cCRE scores. Evaluation of such 39 genes with multi-way interactions revealed that SNPs within multi-way interactions exhibited significantly stronger SNP-SNP interaction effects than expected by chance. Specifically, the SNP-SNP interaction F-test − log_10_(p-values) of the 321 (Gene, SNP_1_, SNP_2_) tuples were notably greater overall than those obtained from permuting the gene expression levels (**Fig**. 3c). A striking example is observed for the gene DKK3, which is linked to Amyloid-*β* pathology and synapse restoration ^34^ (**Fig**. S20, **Fig**. 3d,e). MINTsC identified a multi-way interaction involving ± 200bp of the DKK3 TSS and SNPs rs7480026 and rs16910272 in the 3rd and 5th introns of DKK3, respectively. Statistical testing in a linear model of DKK3 expression revealed a significant interaction effect for the DKK3 expression (FDR≤0.05). Specifically, this analysis highlighted that while each SNP individually has a weak effect, their interaction significantly contributes to DKK3 expression variation. A similar epistatic effect was revealed for other genes (**Tab**. S5), including CPLX2, linked to cognitive resilience and synaptic plasticity ^35^ (**Fig**. S21). These results underscore MINTsC’s potential to discover epistatic effects in molecular QTL studies by generating candidate interactions to test for and, thereby, significantly reducing the multiple testing burden.

## 3 Discussion

We presented MINTsC as an empirical Bayes framework to infer multi-way interactions from pairwise interaction signals quantified in scHi-C data. Despite the limitations due to data sparsity and proximity ligation, scHi-C data harbors cliques representing potential multi-way interactions (albeit of lower sizes ≤ 6) and presents an exciting opportunity to learn higher-order chromatin interactions. MINTsC builds on a dirichlet-multinomial model that takes into account inherent genomic distance bias of scHi-C data by a natural spline model. We developed a computationally efficient estimation strategy for MINTsC and showed that null generative model of MINTsC results in well calibrated p-values, enabling appropriate false discovery rate control (**Fig**. S22, S23, **S24, S25, S26)**. Evaluations of MINTsC using a wide range of real scHi-C datasets yielded multiple-levels of support for MINTsC multi-way interactions from independent external genomic data resources. In addition, multi-way interactions involving genes and their enhancers were strongly supported by their corresponding ABC scores of gene regulation. This finding highlights the potential of MINTsC to infer multi-way gene regulation. Despite the lack of computational methods aimed at identifying multi-way interactions from scHi-C data, we considered a baseline approach by utilizing state-of-the art scHi-C loop calling tool SnapHi-C^11^ and generated multi-way interactions as cliques from significant loop calls. Applications of the baseline approach and MINTsC on scHi-C from human prefrontal cortex cells and synthetic data mimicking actual data yielded that MINTsC is able to identify more multi-way interactions with markedly higher reliability in terms of FDR control. As biological discovery potential of MINTsC, we showcased that MINTsC-derived multi-way interactions in human prefrontal cortex enable discovery of potential epistatic SNP effects. Striking examples included strong SNP-SNP interactions on the expression levels of the genes related to Amyloid-*β* pathology (DKK3) and synaptic plasticity (CPLX2) based on the ROS/MAP study of Alzheimer’s disease. We expect that with the continued development of scHi-C technologies, MINTsC analysis of scHi-C data will complement data from other multi-way interaction assays.

While MINTsC applications presented here relied on known cell type labels, this is not an inherent limitation because there is a plethora of computational tools for inferring cell type labels from scHi-C data ^11, 36–42^. Furthermore, once we quantify the uncertainty of label assignments (e.g., by soft-threshold in clustering of low-dimensional embeddings of single cell data), these uncertainity estimates can be naturally incorporated into the generalized linear model fit that MINTsC employs. More generally, from a methodological point of view, the two-stage (band level and locus pair level) modeling by MINTsC offers a computationally useful framework for any type of high-dimensional and sparse single cell genomics/epigenomics data where exponential family of distributions is a valid generative distribution choice. Specifically, if a group of genomic loci (or genes) can be bundled into a larger class (analogous to a band in MINTsC) for a GLM fitting, then the original locus level estimation (analogous to locus pair level estimation in MINTsC) can be carried out based on estimate allocation procedure introduced in Fig. 1a.

## Acknowledgments

This work was supported by National Institutes of Health (NIH) grants (HG003747, HG012881) and a Chan Zuckerberg Initiative (CZI) single-cell data insights grant (cycle # 1). We thank Dr. Longzhi Tan from Stanford University for insightful discussions regarding the method validation, Dr. Jiansen Lu from Peking University for sharing detailed information and discussions about the scNanoHi-C data, Dr. Takeo Narita from Copenhagen University for sharing the ABC scores of the mESC system, and Siqi Shen from the University of Wisconsin-Madison for sharing processed Nagano *et al*. scHi-C data. We also thank Dr. Ye Zheng from the University of Texas MD Anderson Cancer Center for a useful discussion regarding the baseline method comparison. The results published here are in whole or in part based on data obtained from the AD Knowledge Portal (https://adknowledgeportal.org). Study data were provided by the Rush Alzheimer’s Disease Center, Rush University Medical Center, Chicago. Data collection was supported through funding by NIA grants P30AG10161 (ROS), R01AG15819 (ROSMAP; genomics and RNAseq), R01AG17917 (MAP), R01AG30146, R01AG36042 (5hC methylation, ATAC-seq), RC2AG036547 (H3K9Ac), R01AG36836 (RNAseq), R01AG48015 (monocyte RNAseq) RF1AG57473 (single nucleus RNAseq), U01AG32984 (genomic and whole exome sequencing), U01AG46152 (ROSMAP AMP-AD, targeted proteomics), U01AG46161(TMT proteomics), U01AG61356 (whole genome sequencing, targeted proteomics, ROSMAP AMP-AD), the Illinois Department of Public Health (ROSMAP), and the Translational Genomics Research Institute (genomic). Additional phenotypic data can be requested at www.radc.rush.edu.

## Methods

### MINTsC model representation

MINTsC, presented in **Fig**. 1, is an empirical Bayes model for inferring multi-way interactions from scHi-C contact matrices. It leverages pairwise interactions quantified in contact matrices to construct a test statistic for each candidate multi-way interaction, i.e., clique of bins/genomic-intervals, and evaluates these against a generative null distribution for contact matrices. Here, the candidate multi-way interactions are the cliques whose nodes (loci) are simultaneously observed in at least certain number of cells in scHi-C contact matrices. This step is designed to rule out false cliques that can be detected by merely aggregating pairwise interactions observed across different groups of cells. Specifically, a clique finding algorithm was deployed on each individual cell’s contact matrix. Then, the union of set of these cliques were filtered to retain the cliques that were observed in at least *c* cells across all the cell types. Different *c* values (*c*_0_ and *c*_1_) were employed for 3-way cliques and higher order cliques (with size larger than 4), due to the overwhelming number of size 3 cliques. Consequently, for each cell type, the cliques that were not fully observed (with all of its pairwise interactions) at least once in any of the cells within that cell type were filtered. We defer further details on the choice of these parameters to the Supplementary Note.

For the evaluation of the pre-defined candidate multi-way interactions, we specifically consider cell *k, k* ∈ [*K*], and its symmetric (*cis*-interaction) contact matrices (maps) with dimensions *l*_*chr*_ × *l*_*chr*_ for each chromosome *chr* ∈ [*Chr*], where *l*_*chr*_ denotes the number of loci for chromosome *chr* and the bracket notation [*K*] denotes a set of sequential numbers {1, ·· ·, *K*} (**Fig**. 1). Each entry (edge) of a contact matrix (graph) quantifies the level of contact (edge weight) between two genomic loci (nodes), e.g., *v*_1_ ∈ [*l*_*chr*_], *v*_2_ ∈ [*l*_*chr*_]. None-zero pairwise contacts among more than two loci, e.g., a size 3 clique of loci *v*_1_, *v*_2_, *v*_3_ (starred entries in **Fig**. 1c), provide evidence for a potential multi-way interaction among the loci *v*_1_, *v*_2_, *v*_3_ (triangles in **Fig**. 1c) within a single cell. Under a noise free setting, all pairwise interactions stemming from a multi-way interaction would be observed and generate a clique. However, owing to sparsity levels of contact matrices (different edge widths in each triangle in **Fig**. 1c) and potential proximity ligation limitations, all pairwise interactions may not be observable within an individual cell. To address this issue for detecting multi-way interactions from contact matrices, we view each contact matrix of a cell as an independent sample of the true unknown contact matrix of a context, e.g., cell type, pseudo-time, or treatment. This inherently requires that the cells, to which the method is applied, should be homogeneous enough (Supplementary Note on the homogeneity assumption). Therefore, a user needs to first start with identifying a group of relatively homogeneous cells and apply MINTsC for each group. Then, we compute a clique level score (z-score and p-score) for each candidate clique *q* ∈ [*Q*], by summarizing the pairwise interaction levels of the clique throughout the cells (box with order statistics in **Fig**. 1b). This score aggregates the evidence for individual cliques across the cells of the same type by leveraging all pairwise interactions.

To assess the significance of the cliques, we evaluate whether the clique scores are higher than expected under a contact matrix generating model which considers counts in each upper triangular entry of the symmetric contact matrix of a cell *k* as a random draw from a multinomial distribution with the sequencing depth *C*_*k*_ of the cell as the number of trials. In order to take into account the well-known genomic distance bias, i.e., band bias indicating that contact counts between two loci systematically vary with the genomic distance between them, observed in both the bulk and single cell Hi-C data ^39, 44, 45^, we model the parameters of the multinomial distribution with a natural cubic spline (**Fig**. 1b). Specifically, we define the set of locus pairs within each off-diagonal of the contact matrix as a “band” (gray and gold bars in **Fig**. 1c), where the locus pairs *i* ∈ [*N*_*j*_] in the same band *j* ∈ [*J*] have the same genomic distance *d*_*j*_.

Then, the contact counts *y*_*ijk*_ of locus pairs *i* ∈ [*N*_*j*_] of bands *j* ∈ [*J*] of cell *k* ∈ [*K*] follow a multinomial random vector 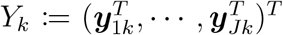 with total count *C*_*k*_, where 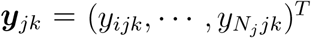. Without loss of generality, we dropped the chromosome subscript *chr* ∈ [*Chr*] and cell type subscript *t* ∈ [*T* ] for multinomial sample *Y*_*k*_; however, this model applies for each chromosome and for each cell type, separately. Additionally, while the contact counts from different cells can be assumed to be independent, dependency among contact counts of a clique within a single cell is plausible. This aspect is naturally incorporated by the multinomial model.

In addition to the multinomial likelihood that serves to estimate the band-level parameters *p*(*d*_*j*_) (Band level Poisson Spline in **Fig**. 1a), we further impose a Dirichlet prior distribution (Gray arrow in **Fig**. 1a) on the locus-pair level contact probabilities 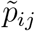. Hyper-parameters *α*_*ij*_ of this Dirichlet prior are utilized to distribute the band-level contact probability *p*_*j*_ onto locus pairs within the band while taking into account the overall null hypothesis (Locus pair level null allocation in **Fig**. 1a). This statistical model for the scHi-C contact counts, along with the null distributional assumption introduced in the next section, enables constructing clique level test statistics, including clique z-score and clique p-score, which enable false discovery rate (FDR) control (**Fig**. 1b, c).

#### Details on statistical framework and estimation of MINTsC

MINTsC relies on a Dirichlet-multinomial distribution that is fitted for each chromosome and cell type separately. In the presentation below, without loss of generality, we drop the chromosome and cell type index. In the MINTsC model, for each locus pair *i* ∈ [*N*_*j*_] within band *j* ∈ [*J*] and a cell *k* ∈ *K*, pairwise contact levels *y*_*ijk*_ follow a multinomial distribution

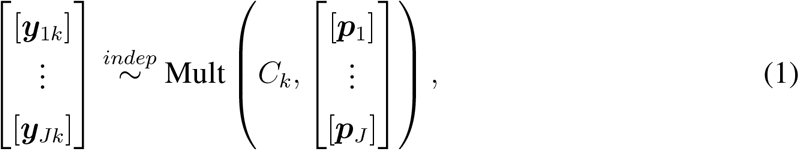

where 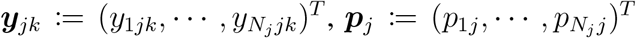, and the number of trials *C*_*k*_ is the sequencing depth of cell *k*. The bracket [·] for each ***y***_*jk*_ (or ***p***_*j*_) emphasizes that this component is vector-valued for each band *j* ∈ [*J*]. We further parametrize the contact probabilities *p*_*ij*_ as 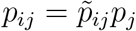, where for 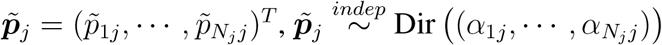 . This formulation includes a band-level probability *p*_*j*_ and within band allocation weights 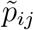. Since 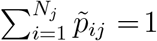 1, by the aggregation property of multinomial distribution, we have the following band-level model

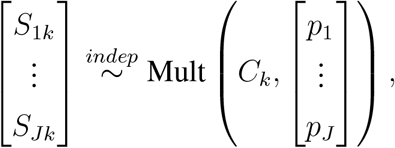

where 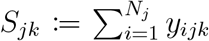 and 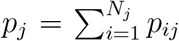 for ∀*j* ∈ [*J*],*k* ∈ [*K*]. Moreover, by the Poisson transformation of the multinomial distribution ^46^, we equivalently have

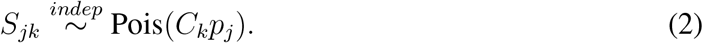

In order to incorporate the well-established band bias (or genomic distance bias) of chromatin conformation capture assays, we further impose a spline model on the band-level contact probability as *p*_*j*_ = *p*(*d*_*j*_), where *d*_*j*_ is the genomic distance between the loci in a locus pair in band *j*. A similar distance dependence was adapted for loop identification in bulk Hi-C data ^44^. Here, *p* is a natural-cubic spline function with *L* number of basis functions and is set to *L* = 20, 20, 30, 30, 100 for Ramani *et al*., Li *et al*., Nagano *et al*., Tan *et al*. and Lee *et al*. data, respectively, in the applications presented.

MINTsC first fits the model in **Eqn**. (2), i.e., obtains 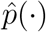, by a Poisson generalized linear model on the total band counts *S*_*jk*_ (Band level Poisson Spline in **Fig**. 1a). This procedure reduces computational burden significantly (e.g., from a spline GLM fit with the number of experimental units as {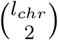 locus pairs ∀*chr* ∈ [*Chr*] to only {number of bands}, with as many observations as the number of cells for the cell type for each experimental unit. Then, within each band *j*, MINTsC distributes the estimated overall band contact probability 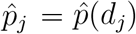 across all the locus pairs (Locus pair level null allocation in **Fig**. 1a) under the following null assumption:

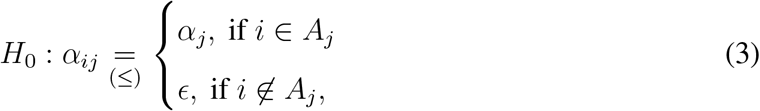

where *ϵ >* 0 is a small positive number, e.g., *ϵ* = 1*e* − 05. Here, *A*_*j*_ is the set of band *j* locus pairs that are expected to have nonzero contact counts, and, similarly, 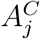 is the set of locus pairs that are more likely to have zero contacts. This null assumption results in expected contact counts as 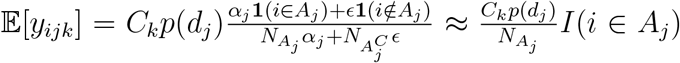, where 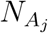 denotes the cardinality of the set *A*_*j*_. This intuitively indicates that the locus pairs in *A*_*j*_ will share the non-zero uniform expected contact counts *C*_*k*_*p*(*d*_*j*_), and the ones outside of *A*_*j*_ will have expected count as zero. The set *A*_*j*_ can be obtained from external data including combination of mappability scores and GC content. However, given that we are in a setting where we are considering multiple cell types, these sets can be obtained by evaluating the contact counts of the loci across all the cells excluding the cells from the cell type that is being analyzed (Supplementary Note for details on the construction of *A*_*j*_ set). This exclusion is to ensure that this set, which informs the null assumption, is estimated from independent data. A simpler choice for *α*_*ij*_ would be equal distribution of the contact probabilities across all the locus pairs within a band: 1*/N*_*j*_. However, our experiments indicated that this choice of prior null underestimates the mean of the clique statistics and leads to uncalibrated p-values as quantified by significant violations from standard normal in the resulting Q-Q plots. We discuss how *A*_*j*_ can be set by leveraging cells of the other cell types in the next section.

Given the credibly non-zero set *A*_*j*_, we can estimate *ε*_*ij*_ with an empirical Bayes approach by maximizing the marginal log likelihood 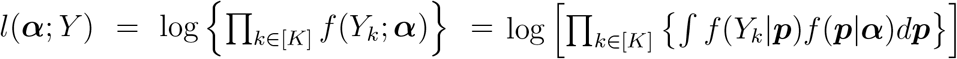 as:

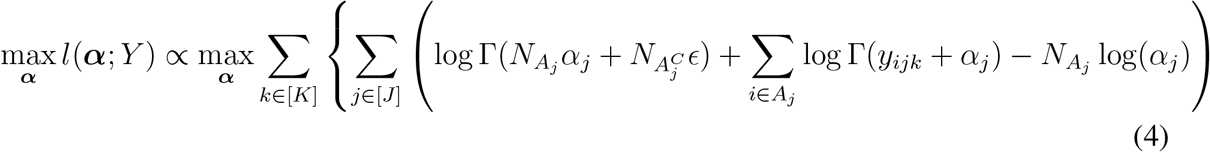

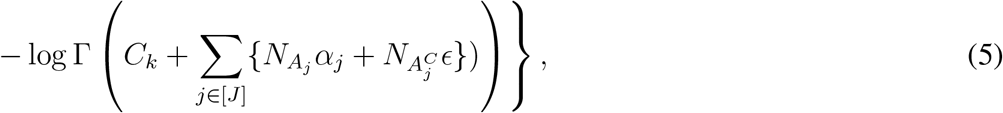

under the null hypothesis **Eqn**. (3) on *α*, where 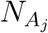(or 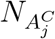 ) denotes the cardinality of the *A*_*j*_ (or 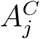) set, *Y* = (*Y*_1_, ·· ·, *Y*_*K*_), *Y*_*k*_ = (***y***_1*k*_, ·· ·, ***y***_*Jk*_), 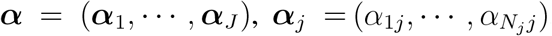, and *p* = (*p*_1_, ·· ·, *p*_J_ ). Then, using the properties of conditional expectation and covariance, under the null model **Eqn**. (3), we have locus pair level null parameters

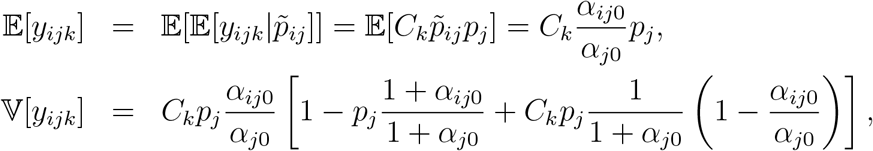

where 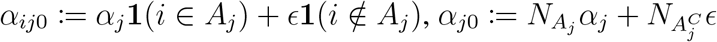. For each locus pair *i* (within band *j*) within a size *N* clique *q* for *m* ∈ [*M* ], where 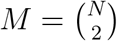, we define

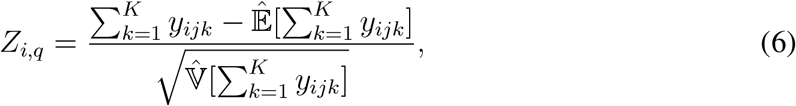

where both the expectation and the variance terms are estimated under the null assumption on the locus pair. As the random variable 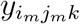 is independently distributed across cells *k* ∈ [*K*], by the Lindeberg-feller Central Limit Theorem and the 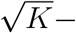consistency of the Maximum Likelihood Estimator (MLE), under some regularity conditions required for MLE, we achieve

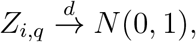

as the number of cells *K* → ∞ under the null distribution assumption **Eqn**. (3).

Finally, based on the set of locus pair level statistics of a (size *N* ) clique *q*, we introduce clique level summary statistics. First, considering the vector of locus pair level statistics within a clique *q*, ***Z***_*q*_ = (*Z*_1,*q*_, *Z*_2,*q*_, ·· ·, *Z*_*M,q*_) with 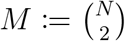, we define the *r*th largest value *Z*_(*M≥r*+1)*q*_ as the *clique z-score* of the clique *q*. Equivalently, for ***p***_*q*_ = (*p*_1,*q*_, *p*_2,*q*_, ·· ·, *p*_*M,q*_), where *p*_*i,q*_ := 1 − Φ (*Z*_*i,q*_) denotes the right-tail standard normal cdf value of *Z*_*i,q*_, we let the *r*th smallest value *p*_(*r*),*q*_ as the *clique p-score* of the clique *q*. Hence, the clique level summary statistics are defined as follows for clique *q*:

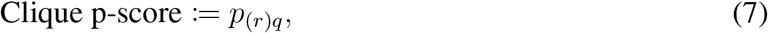

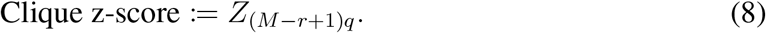

The key reasoning behind choosing *r*th strongest locus pair level statistics of a clique is that we require a clique to have at least *r* number of significant pairwise edges to represent a multiway interaction. A small enough *p*_(*r*)*q*_ (or large enough *Z*_(*r*)*q*_ ) automatically guarantees that *r* number of locus pairs will have significant interactions to form a multi-way interaction. Under this definition, we have tractable null distribution for the clique p-scores. Theoretically, for independent *p*_*iq*_s following uniform distribution *U* (0, 1), we have *p*_(*r*),*q*_ ∼ Beta(*r, M* − *r* + 1). We note that the entries of ***Z***_*q*_, and therefore those of ***p***_*q*_, are not necessarily independent of each other due to the multinomial assumption. However, empirically, the covariance between the entries are sufficiently small (with ratio approximately 1*/*100) compared to the variance terms across all the datasets we analyzed. Therefore, we approximate the null distribution of the clique p-score as

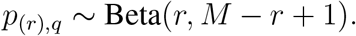

Analytically tractable p-values of the p-scores facilitate false discovery rate control in the downstream analysis. Clique-level p-values are computed from the parametric distribution Beta(*r, M* − *r* + 1), i.e.,

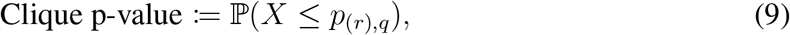

where *X* ∼ Beta(*r, M* − *r* + 1). Then, the Benjamini-Hochberg procedure is employed with these p-values for FDR control across the candidate cliques.

The most conservative choice for *r* is *M*, where the maximum of the locus pair level p-values of a clique is set as the clique p-score. Empirically, we (and others in a different context for combining p-values ^47^) observed that when the observed group of p-values, e.g., a clique’s locus pair level p-values in this case, exhibit deviations from the assumed theoretical null due to data noise, maximum p-value or a too large choice of *r* is not robust, i.e., its empirical null distribution deviates from the theoretical null distribution, especially for large *M* ^47^. Moreover, in practice, this leads to detecting too few cliques. Another choice of *r*, 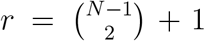, ensures that all the *N* loci participate in forming a connected graph, when *p*_(*r*)*q*_ is small enough. However, our empirical results show that this choice is not well calibrated when compared to Beta(*r, M* −*r* +1). Hence, for the datasets analyzed in this paper, we set *r* = *N* −1 for a size *N* clique and filter the cliques from further testing if the *r* strongest locus pairs involve fewer than *N* loci in order to make sure the significant pairwise edges at least form a connected graph. The latter condition ensures that all the loci within the clique are involved in at least one pairwise interaction. The order statistic summarization to construct a clique score naturally makes the multi-way interactions to involve a sufficient number of strong pairwise interactions, while the non significant cliques would involve a relatively smaller number of significant pairwise interactions.

#### Details of synthetic data generation

We now introduce details on synthetic data generation for the analyses made in Section 2.2 and 2.6.

For both of the simulations, we leveraged Lee *et al*. ^22^ scHi-C dataset (chromosome 19 with 5,582 bins of 10Kb, and L2/3 neurons) and the model in **Eqn**. (1), where the multinomial contact probabilities 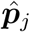, *j* ∈ [*J*] were initially estimated under the null assumption **Eqn**. (3). Then, in the null contact matrices learned from Lee *et al*. ^22^ scHi-C data, we implanted clique structures for the simulation-based analyses.

For the analysis made in Section 2.2, in order to showcase how MINTsC empirically handles spurious cliques that are mere aggregation of pairwise locus pairs appearing in disjoint set of cells, two different sets of clique structures were implanted, which resulted in the population level contact probability matrix as a mixture of two contact matrices. Specifically, both matrices had the same 300 three way cliques (with edges *A*_*i*_, *B*_*i*_, *C*_*i*_, *i* = 1, ·· ·, 300) which constituted the true three-way interactions. For the other 300 three-way cliques (referred to as spurious multi-way interactions) with edges (*D*_*i*_, *E*_*i*_, *F*_*i*_, *i* = 1, ·· ·, 300), one of the matrices had the *D*_*i*_ and *E*_*i*_ edges, whereas the other had the *F*_*i*_ edge to ensure that two of the three edges were observed in half of the cells, while the third edge was observed in the other half. This mimics the setting where aggregated pairwise data might spuriously suggest a three-way interaction (**Fig**. S5a). Then, cells were divided into two groups, and contact counts for each cell were generated by sampling from one of the two contact matrices, with the first half of the cells sampled from the first matrix and the remaining half from the second. In both cases, contact counts were sampled from a multinomial distribution with probabilities given by the corresponding contact matrix. Across 551 simulated cells in one simulation replicate, each true three-way interaction was observed in an average of 7.4 (median of 7) cells and each spurious three-way interaction was observed in an average of 1.7 (median of 0) cells. To provide practical validity of the simulated true cliques, we analyzed a CRISPRi validated multi-way interaction involving the MYC gene in K562 cells^1^. We quantified how many cells in the K562 scHi-C dataset contained the complete clique corresponding to this multi-way interaction. Out of 135 cells, only 7 cells (5%) harbored the full set of pairwise contacts, which roughly mimics the observation we have for the true cliques. All true multi-way interactions were observed as cliques in at least one cell, while only 11% of the spurious multi-way interactions were observed as cliques in at least one cell. These patterns are consistent with our empirical findings from multiple scHi-C datasets, where the number of cells containing cliques tends to be limited due to sparsity of the data. Specifically, in any given scHi-C dataset, only about 2–5% of the cliques are observed in more than 10 cells.

With the generated data, we assessed MINTsC’s ability to distinguish between these two groups across varying levels of interaction signal (i.e., contact probabilities in the population level contact matrices) and ten replicated simulations. With moderate signal strength (**Fig**. S5b), MINTsC achieved an 83% true positive rate (TPR) and an 8% false positive rate (FPR). With strong signal, it achieved 100% TPR and 6% FPR, consistent with the expected behavior under a 5% FDR threshold (**Fig**. S5c).

For the baseline comparisons made in Section 2.6, to ensure a more heterogeneous background contact count distribution than accounted by MINTsC (i.e., not strictly following the MINTsC background), we considered 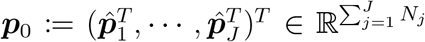, where the multinomial contact probabilities 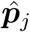, *j* ∈ [*J*], of model in **Eqn**. (1) were estimated under the null assumption **Eqn**. (3), and introduced heterogeneity through 2,100 background cliques with pairwise interaction signal size of 0.00001. As the signal contact counts, we considered 300 multi-way interaction cliques with pairwise interaction signal sizes of 0.000025 (Lowest signal), 0.00005 (Low), 0.000075 (Medium), and 0.0001 (High). This construction resulted in a parameter vector ***p***_*a*_, each entry of which denoted (population) pairwise interaction value of the implanted background (2,100) and true (300) cliques. Then, we generated the scHi-C contact maps across cells *k* ∈ [*K*] based on

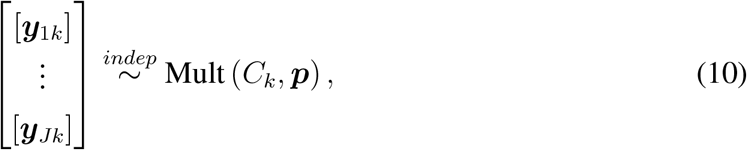

where ***p*** ∝ ***p***_0_ + ***p***_*a*_. Specifically, in each simulation replicate, we generated contact maps across 551 cells under each signal setting (Lowest, Low, Medium, High) and kept the number of simulation replications at 10 to ensure computational feasibility of the baseline method (Supplementary Note).

Within each simulation replicate across the four signal settings, we applied MINTsC to infer multi-way interactions (at FDR 0.05) and also fit SnapHi-C, which is designed to exclusively call loops, for the baseline comparison. Significant SnapHi-C loops (at FDR of 0.05) were aggregated into cliques to identify multi-way interactions. Inferred multi-way interactions from both approaches were evaluated for their performance in identifying the true and false multi-way interactions.

